# Avifauna recovers faster in areas less accessible to trapping in regenerating tropical forests

**DOI:** 10.1101/2021.08.20.457106

**Authors:** H S Sathya Chandra Sagar, James J. Gilroy, Tom Swinfield, Ding Li Yong, Elva Gemita, Zuzana Burivalova, Novriyanti Novriyanti, David C. Lee, Muhammad Nazri Janra, Andrew Balmford, Fangyuan Hua

## Abstract

Tropical forest restoration stands to deliver important conservation gains, particularly in lowland Southeast Asia, which has suffered some of the world’s highest rates of forest loss and degradation. This promise, however, depends on the extent to which biodiversity at forest restoration sites continues to be exposed to threats. A key knowledge gap concerns the extent to which biodiversity recovery in naturally regenerating tropical forests is impacted by trapping for the multi-million-dollar wildlife trade. Here, we use a repeated survey dataset to quantify rates of avian community recovery under forest regeneration, at a flagship restoration site in the lowland rainforests of Sumatra, Indonesia. We show that over a decade, forest regeneration was associated with significant abundance increases for 43.8% of bird species.

However, the apparent negative impacts of trade-driven trapping on avian populations also intensified: the proportion of species that show increased abundance only in very remote forests increased from 5.4% to 16.2%, while the overall accessibility of the forest increased. We found that 14% of species did not recover as fast as predicted based on the observed improvement in forest conditions over the study period. Our results highlight the potential for rapid avifaunal recovery in regenerating tropical forests, but also emphasize the urgency of tackling the serious threat of wildlife trade to Southeast Asia’s biodiversity.

## 1. INTRODUCTION

Tropical forests worldwide have undergone widespread loss and degradation with severe consequences for biodiversity, people, and critical ecosystem services (Barlow et al., 2018; Edwards et al., 2019; Gibson et al., 2011; Watson et al., 2018). While the protection of existing old-growth forests is paramount, restoration of degraded lands can also deliver important conservation gains (Chazdon and Brancalion, 2019; Lewis et al., 2019; Strassburg et al., 2020; Watson et al., 2018). This is particularly true for Southeast Asia, where only 8.4% of the historical old-growth forest remains intact (Potapov et al., 2017; Sodhi et al., 2010; Wilcove et al., 2013). The future of biodiversity in this region, among the richest in the world, depends to a large extent on the effective and timely restoration of its forest habitats (Cosset and Edwards, 2017; Edwards et al., 2014, 2009; Senior et al., 2019). While the increasing momentum of forest restoration in Southeast Asia is encouraging (Chazdon et al., 2017; FAO and UNEP, 2020), the realization of its conservation success hinges on tackling the negative biodiversity impacts of other threats in forests undergoing restoration, especially wildlife trapping.

Across Southeast Asia, wild bird trapping driven by the pet trade poses a severe conservation threat (J. A. Eaton et al., 2015; Harris et al., 2017; Symes et al., 2018). The pet bird trade in the region, part of a global issue, is estimated to be worth hundreds of millions of dollars annually (Hughes, 2021; Marshall et al., 2019; Morton et al., 2021). It affects thousands of species, particularly those targeted for singing competitions and pet-keeping (Jepson, 2010; Scheffers et al., 2019; Shepherd, 2006). Market and household surveys in Indonesia suggest that the pet bird trade is ubiquitous across the country and that most traded birds are sourced illegally from the wild (Burivalova et al., 2017; Chng et al., 2015, 2018a; Shepherd et al., 2004). There is evidence that the scale of trade has increased over the past decade driven in part by increased accessibility due to forest loss and degradation (Marshall et al., 2019). Limited field evidence has linked increase in trapping, to decrease in bird populations in the wild (Harris et al., 2017). Together, this suggests that trade-driven trapping could dampen the recovery of bird populations in forests undergoing restoration in Southeast Asia.

In this study, we evaluated the recovery of avian diversity over 10 years of forest restoration in a region increasingly impacted by trade-driven trapping, at a flagship ecosystem restoration site in the now heavily modified lowlands of Sumatra, Indonesia. We conducted repeated bird surveys at the community level, sampling across gradients of forest condition and trapping pressure. We examined how species abundance changed over time, its relationship with forest conditions and trapping pressure, and the extent to which its recovery had been affected by intensifying trapping pressure. We also assessed how species recovery related to the market demand, habitat association, and IUCN Red List status of each species.

## 2. METHODS

### 2.1 Study site

We conducted our study in the Harapan Rainforest (‘Harapan’ hereafter), which straddles the provinces of Jambi and South Sumatra in Sumatra, Indonesia (2°08′ S, 103°22′ E, 50–80 m a.s.l.; Fig. 1). Harapan was established as Indonesia’s first ecosystem restoration concession in 2007. It was jointly managed by a consortium of conservation organizations with heavy financial investment and on-the-ground presence since 2008, with the main goal of recovering biodiversity in forests after logging (Harrison and Swinfield, 2015; Hua et al., 2016; Lee et al., 2019, 2014). As of 2018, it covered 98 555 ha of lowland dipterocarp forests in various stages of recovery, after commercial selective logging ceased in 2005. Rich in biodiversity representative of the Sundaic lowlands, it is also recognized as an Important Bird and Biodiversity Area (IBA; BirdLife International, 2017). Since its establishment, Harapan has faced mounting conservation challenges: as of 2018, it lost ∼25 000 ha of forest cover to illegal logging and encroachment, and another ∼30 000 ha was damaged by El Niño-related drought and fires in 2015 (Fig. S1), particularly the eastern and the northern regions. Despite these challenges, Harapan’s remaining forests stayed contiguous and largely demonstrated signs of recovery through natural regeneration, and at some heavily degraded sites through active tree planting (Fig. 1).

**Figure 1:**
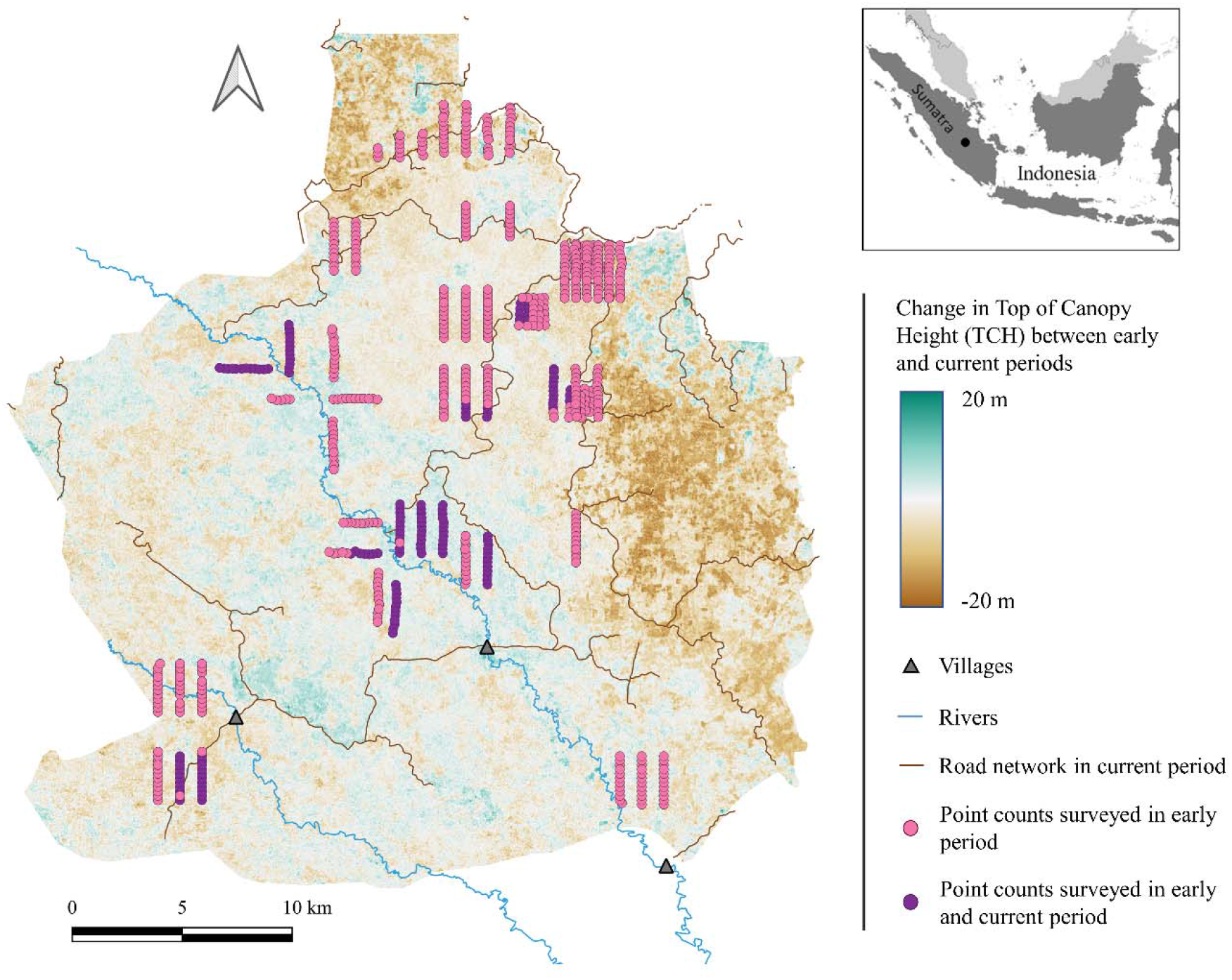
Location of point count stations and changes in top-of-canopy height (TCH) over time at Harapan Rainforest. Point count stations were surveyed either during the early period (2009-2011, pink); or during both the early and current periods (2009-2011, 2018, purple). Change in TCH was estimated from Landsat imagery using a LiDAR training dataset.

Wildlife trapping and hunting, especially of birds, has been persistent in Harapan (pers. comm. with local bird trappers and Harapan staff, 2018), aided by a network of seasonally navigable roads and rivers that allowed access to most places within the area (Fig. 1), and the lack of anti-poaching patrols except for those targeting a few charismatic mammal species. Anecdotal evidence indicated that bird trapping in Harapan had intensified in the years leading up to our study, likely linked to increasing human accessibility (pers. comm. with Harapan staff, 2018). Trappers used various methods to capture live birds for the pet market, including mist nets, live bird traps, and strong glue. In recent years, the use of shotguns in Harapan to kill hornbills had also been recorded (pers. comm. with bushmeat hunters, 2018), often targeting the Critically Endangered Helmeted Hornbill *Rhinoplax vigil* that is prized for its ivory-like casque (Beastall et al., 2016; BirdLife International, 2020).

### 2.2 Bird community surveys

We surveyed bird communities in Harapan in 2009-2011 (‘early period’ hereafter; Lee and Lindsell 2011; Hua et al. 2016) and again in 2018 (‘current period’ hereafter) using point counts. We positioned point count stations ≥200 m apart so that they are spatially independent of each other, along line transects that covered a range of forest conditions and human accessibility (as a proxy for trapping pressure; see section 2.4). In total, we surveyed 636 stations in the early period (Fig. 1). Of these stations, only 287 stations were in the contiguous forest portions of Harapan that were spared from significant fire and deforestation between the early and current periods (Fig. S1), and we confirmed the recovery of their forest habitat via remote sensing analysis (see section 2.3). We focused on these stations for the current survey, from which we selected 144 stations that covered a range of forest conditions and human accessibility using a stratified random sampling design (Fig. 1).

For both early and current periods, we conducted unlimited radius point counts for the entire bird community that allowed for the correction of imperfect detection in estimating species abundance, excluding nocturnal, wetland, aerial, or raptorial species. We employed 10-minute counts in the early period and 12-minute counts in the current period. Given our correction for imperfect detection in data analysis (see section 2.6), the different lengths of point count between the early and current periods should not bias our estimation of species abundance. During each point count, we recorded all birds seen or heard (excluding flyovers), along with their time of initial detection in minutes since the onset of the point count period. The latter information allowed us to use removal models to estimate and correct for species’ detection probability (Farnsworth et al., 2002). We conducted all surveys between 05:30 – 11:30 on days without rain or strong wind, and we recorded the time of the onset of each point count in minutes since sunrise (‘survey time’ hereafter). The early-period surveys were conducted by Harapan’s research team led by DCL (551 point count stations; Lee and Lindsell, 2011) and by FH in a separate research project (85 point count stations; Hua et al., 2016), while surveys for the current period were conducted by HSSCS (144 point count stations).

We took two measures to minimize the potential bias in bird community characterization due to different observers conducting surveys. First, we used observer identity as a random effect for detection probability in our removal models, thereby correcting for potential detection differences among observers in estimating species abundance (see section 2.6). Second, considering that the varying survey skills of the different members of Harapan’s research team may bias the ability to detect some small (e.g., White-chested Babbler, *Pellorneum rostratum*) or inconspicuous (e.g., Grey-chested Jungle-Flycatcher, *Cyornis umbratilis*) species, for these ‘prone-to-miss’ species, we used only the subset of data collected by FH for the early period. We identified these species by assessing the number of times a species was detected out of the pool of point count stations (‘detection rate’ for short) between the subset of data collected by Harapan’s research team *versus* by FH: we considered a species as prone-to-miss if its detection rate in the former sub-dataset was ≤20% of that in the latter sub-dataset (Table S1).

### 2.3 Quantifying forest condition

We measured changes in forest condition across the study periods using the metric of top-of-canopy height (‘TCH’ hereafter). We estimated TCH from a time series of Landsat imagery, using a model derived from LiDAR training data through a machine learning approach. The LiDAR images were collected by TS for a different research project on October 24, 2014, that covered 3 626 ha (3.7%) of our survey area (Fig. S2). We processed the LiDAR data to a 0.5-m-resolution TCH model as described in Swinfield et al. (2019), and aggregated and resampled the TCH values to a 30-m resolution to align with Landsat imagery. Next, we used all Landsat imagery covering Harapan within 1 year of each of the two bird surveys and the LiDAR survey to predict TCH (Asner et al., 2018; Csillik et al., 2019). For this purpose, we converted the surface reflectance values of Landsat imagery to five vegetation indices (VIs) considered consistent between remotely sensed scenes and suitable for estimating vegetation height (Appendix A; Jin and Sader 2005; Xue and Su 2017). We used the VIs from two discrete sets of Landsat images around the LiDAR survey that were two years apart (in 2013 and 2015; Hastie et al. 2009) to train a random forest model for predicting TCH over 75% of the LiDAR areal coverage (Appendix A), using package ‘randomForest’ (version 4.6; Breiman and Cutler, 2018) in R (R Development Core Team, 2018). Testing using the remaining 25% of LiDAR data showed good model performance (Fig. S2). We then predicted TCH for the entire survey area using the VIs derived from the 2009 and 2018 Landsat images (i.e., within 1 year of the bird surveys) and the random forest model.

To represent the TCH of a given point count station in a given period, we averaged the predicted TCH values over all the pixels within a 100 m radius of the station from the appropriate time (‘mean TCH’ hereafter). We opted for a remote sensing approach to measure forest condition across both study periods in a standardized way and at a scale appropriate to the habitat of most bird species. Whereas field vegetation survey had not been carried out at a sufficient breadth during the original survey, we were able to use it to ground truth our remote sensing metric. Our TCH metric was correlated with measurements of tree basal area obtained for a subset of point count stations (early period: r_(474)_ = 0.31, p < 0.01 ; current period: r_(132)_ = 0.2, p < 0.05), indicating its utility in representing forest conditions.

### 2.4 Quantifying human accessibility

For each point count station, we estimated its accessibility to humans as a proxy for the trapping pressure it likely was under, with greater accessibility representing stronger trapping pressure (Harris et al., 2017). The use of this proxy for trapping pressure was necessary due to the difficulty of directly measuring trapping activities across large landscapes over two study periods. Most roads and rivers in Harapan were navigable by motorbikes and boats, which allowed relatively easy access from nearby human settlements. Given that these settlements were observed to be trade hubs for wild-caught birds in and around Harapan, we assumed that the primary determinant of human accessibility to a given location in Harapan would be the effort needed to access it on foot. For each point count station, we calculated its Euclidean distance from the nearest ‘easy-access point’ (i.e. roads, rivers or trails) as a measure of the difficulty of human access (‘access difficulty’ hereafter), using the map of Harapan in the package ‘FNN’ (version 1.1.3; Beygelzimer et al. 2019) in R (R Development Core Team, 2018). We assumed that habitat conditions inside the forest and the seasonality of the river water levels did not influence the effort taken to access a particular location.

### 2.5 Species market demand, habitat association, and IUCN Red List status

We classified all bird species recorded in our surveys into two trade guilds that represented the relative market demand for them, based on the most up-to-date market survey data for the region (Chng et al., 2018a, 2018b, 2016, 2015; Leupen et al., 2018; Rentschlar et al., 2018; Shepherd et al., 2016, 2004): (1) targeted species (high demand) – species that are highly prized and in high demand for their singing abilities (e.g. songbirds), ornamental attractiveness (cage birds) or body parts (e.g. helmeted hornbills); and (2) opportunistically trapped species (generic demand) – this includes all other species that are not specifically targeted but nonetheless trapped as ‘bycatch’ and sold in the market whenever possible. Our classification scheme considered all species as in demand in the market, albeit to different extents. We based this scheme on insights from informal interviews with trappers and local conservationists, which suggested that all trapped birds, if still alive, were supplied to the market. Compared with opportunistically trapped species, we expected that the abundance of targeted species would be more prone to the negative impacts of trapping.

Additionally, we classified all bird species recorded in our surveys into two habitat association guilds, based on the Birds of the World database (Billerman et al., 2020): (1) forest-dependent species – species that prefer primary or mature secondary forests; and (2) generalist species – species that are able to survive in or prefer heavily degraded natural forests, plantations, open areas, or human-dominated landscapes. Compared with generalist species, we expected that the abundance of forest-dependent species would increase more markedly over time as the forest condition improved under restoration (Latja et al., 2016; Owen et al., 2020). Finally, for all bird species recorded in our surveys, we recorded their current IUCN Red List categories, along with descriptions of the threats they face (IUCN, 2019). We compiled this set of information to assess the extent to which the current Red List status and conservation threat assessments reflected the threat posed by trapping as indicated by our research. For species prone to the negative impacts of trapping, we expected the current assessments of their conservation threats to recognize trade-driven trapping as a major threat, and their Red List status reflecting this recognition.

### 2.6 Statistical analysis

#### 2.6.1 Estimating species abundance, its change, and its relationship with predictor variables

We used community-level abundance models (Royle, 2004; Yamaura et al., 2012, 2011) under the removal-model framework (Farnsworth et al., 2002) to estimate the abundance of each species during each study period and its relationship with forest condition and access difficulty. We limited our model-building to species observed during both periods to assess changes in their true abundance over time, because for species not observed in a given survey, it was not possible to estimate their true abundance. Thus, of the 187 species we recorded during both study periods (177 and 132 species during the early and current periods, respectively), we retained 122 species that were recorded in both periods for abundance modeling (Table S2).

To account for imperfect detection under the removal-model framework (Farnsworth et al., 2002), we divided each point count into four time intervals (*t*); intervals were 2.5 and 3 min for the early and current periods, respectively. While the difference in interval lengths between the early and current periods meant that detection probability applied to different lengths of time, this should not confound the estimation of species abundance, as the number of intervals were the same between the periods. We then tallied the number of individuals for each species that were newly detected during each interval *t*, which we expressed as Y_*i,j,k-t*_ for species *i* at point count station *j* during study period *k*. Similarly, we used λ_*i,j,k*_ to represent species *i*’s true mean abundance at point count station *j* during study period *k*, i.e. that expected under the forest condition and access difficulty represented by the point count station. We modeled λ_*i,j,k*_ as a linear function of mean TCH and access difficulty on a log link as in Equation 1, after confirming the lack of strong collinearity between these two variables in both survey periods (early period: r_(656)_ = -0.05, p = 0.177; current period: r_(656)_ = 0.06, p = 0.113).

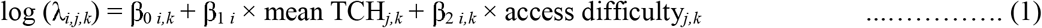

To parameterize Equation 1 in a community-level abundance model (Royle, 2004; Yamaura et al., 2012, 2011), we assumed that species-level coefficients followed a normal distribution that characterized the community-level response; we denoted the mean of the normal-distributions for these community-level coefficients as β_0c *k*_, β_1c_, and β_2c *k*_, respectively, with c denoting the community level. We fixed the coefficient for mean TCH (β_1 *i*_ and β_1c_) across the two study periods, as the response of a species to habitat quality is unlikely to drastically change within around ten years unless there was extreme selection pressure (e.g. Grant et al., 2017). However, we allowed the coefficient for access difficulty (β_2 *i,k*_ and β_2c *k*_) to change across study periods, considering that trapping pressure and its influence on species abundance may have changed over time in Harapan. We modeled the realized abundance of species *i* at point count station *j* during study period *k*, N_*i,j,k*_, as a Poisson draw from the mean λ_*i,j,k*_ as in Equation 2:

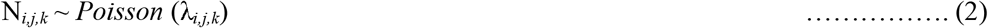

As individual birds were detected (thus ‘removed’) during each successive interval within a point count, we calculated the abundance of birds that remained to be detected during each interval, N_*i,j,k-t*_, as in Equations 3:

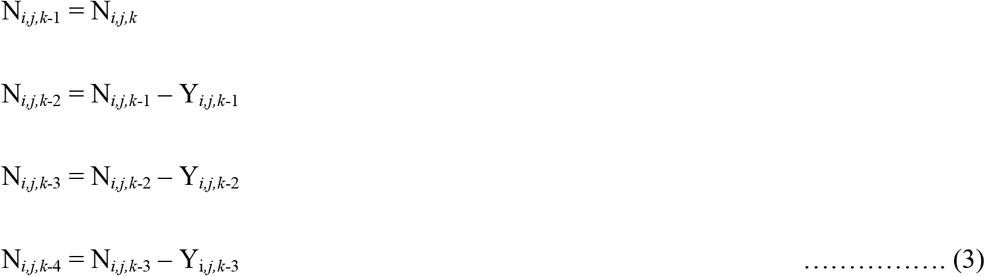

where we modeled the observed count for each interval Y_*i,j,k-t*_ as a binomial variable with N_*i,j,k-t*_ trials and detection probability p_*i,j,k*_, assuming that for species *i* at point count station *j* during study period *k*, this probability was consistent across all intervals, as shown in Equation 4:

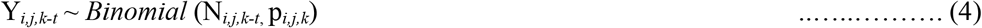

We modeled p_*i,j,k*_ as a linear function of the survey time (i.e. the time at which the point count took place, measured in minutes after sunrise; scaled and centered) on a logit link, treating the identity of observer *m* as a random effect (we considered all members of Harapan’s research team as one observer), as in Equation 5. We assumed a linear relationship between survey time and detection of birds to reduce the risk of overparameterization (Fig. S3).

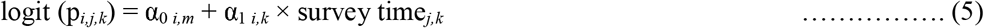

We fitted all models in a Bayesian framework using JAGS (Plummer, 2003) with the packages ‘rjags’ (version 4-10; Plummer 2016) and ‘r2jags’ (version 0.6-1; Su and Yajima 2015) in R (R Development Core Team, 2018). We used uninformative priors, and we ran the model with 100,000 iterations on three chains, with a burn-in of 90,000 and a thinning value of 5. We evaluated the convergence of the model using the Rhat value (mean Rhat of our model = 1.002), which should ideally be close to 1 (Plummer, 2012). Having already a covariate-heavy model, we decided against using a spatial term quantifying landscape configuration in our model to reduce overfitting or parameter identifiability. Additionally, Moran’s I test for spatial autocorrelation among number of species detected across point count station, found a weak autocorrelation in the early period (*Moran’s I* = 0.22, p < 0.01), and for the current period, close to random distribution (*Moran’s I* = 0.07, p < 0.01).

From the posterior distributions of the model, we derived, for each species, and the entire community in each survey period, the median values and 89% Bayesian equal-tailed credible intervals (89% ETI hereafter; Kruschke, 2014; Makowski et al., 2019; McElreath, 2018) for its mean abundance across all point count stations, denoted hereafter as 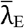 and 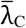 for the early and current periods, respectively. We also derived the median and 89% ETI for the relative change between 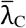 and 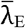 (denoted as 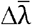 hereafter), and all model coefficients.

#### 2.6.2 Assessing the impacts of trapping pressure on avifaunal recovery

We first assessed the impacts of trapping pressure on avifaunal recovery by assessing the changes over time in model-estimated β_2_ for each species and the entire community, i.e., the relationship between species abundance and access difficulty. Given that a positive β_2_ indicates higher abundance in less accessible areas, likely linked to the negative impacts of trapping, if β_2_ became more positive over time, it would indicate intensified negative impacts of trapping on species abundance. Where we detected β_2_ becoming more positive, we further assessed how intensified trapping impacts may have influenced species abundance recovery as predicted from improved forest conditions. To do this, for each species and the entire community, we used Equation 1 to calculate a counterfactual current mean abundance across all point count stations, had the negative impacts of trapping not intensified (denoted as 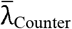 hereafter). For each point count station, we applied its values of TCH and access difficulty to Equation 1, using the coefficients derived from the models above except for β_2_, for which we used the coefficient for the early period. We then calculated the difference between 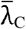 and 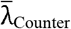 for each species and the entire community; a negative value would represent deficit in its abundance recovery (‘recovery deficit’ hereafter) attributed to the intensification of threats posed by trapping over time.

We further calculated and plotted 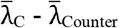 for the entire community across a raster grid of Harapan to visualize the community-level recovery deficit and to identify a ‘deficit zone’ of avian abundance recovery in Harapan, i.e., where 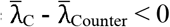, indicating the areas where species’ abundance recovery linked to improvements in forest conditions could not compensate for the decline caused by trapping. We used a similar approach as above, applying Equation 1 while using the TCH and access difficulty values of each grid cell instead of each point count station. We checked post hoc whether the abundance and coefficients derived from our model differed with the trade and habitat association guilds.

## 3. RESULTS

From the 144 point count stations we sampled during both the early and current periods (Fig. 1), we recorded a total of 187 bird species from both periods combined, of which 55 species were detected only during early period and 10 species only during the current period (Table S2). We used 122 species recorded in both periods for abundance modeling (Table S3). Among the species that were only detected in the early period, 76% were opportunistically trapped species in terms of trade guild and 56% habitat generalists in terms of habitat association. Among the species recorded only during the current period, 90% were opportunistically trapped species, with an equal proportion (50%) belonging to forest specialist and habitat generalists.

Over the study period, the TCH metric increased across forest areas in Harapan that did not experience significant fire, degradation, or deforestation (Fig. 1), including at the point count stations resampled (mean difference = 0.38, 95% CI = -0.04, 0.79; paired t-test, t_20_ = 1.8, p = 0.08; Fig. S4). We found that forests in Harapan had become more accessible over time: mean access difficulty and the Euclidean distance of the least accessible point count station from the nearest easy-access point, decreased from 1.32 km (SD = 1.29 km) and 6.71 km respectively in the early period to 0.72 km (SD = 0.57 km and 2.70 km respectively in the current period. We found no relationship between changes in TCH and access difficulty, indicating that improvements in forest condition occurred across the study area, even in areas of high accessibility (Fig. S5).

For the 122 bird species analyzed, 45.1% of species showed significantly greater mean abundance in the current period compared with the early period (i.e., 89% ETI of 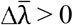), while no species showed significantly lower mean abundance over time (i.e., 89% ETI for 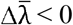; Fig. 2a). At the community level, the mean 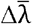 across all species, which represents the mean increase in bird individual numbers for the area covered by a single point count station, was estimated to be 0.69 over early period levels (89% ETI: -0.07% to 1.46; Fig. 2a). With improving forest condition, we found that abundances of 8% of the species increased significantly and decreased significantly for 12.3% of the species. At the community level, we did not detect a significant relationship between species abundance and forest condition (median β_1_: -0.02 with 89% ETI: -0.34 to 0.31).

**Figure 2:**
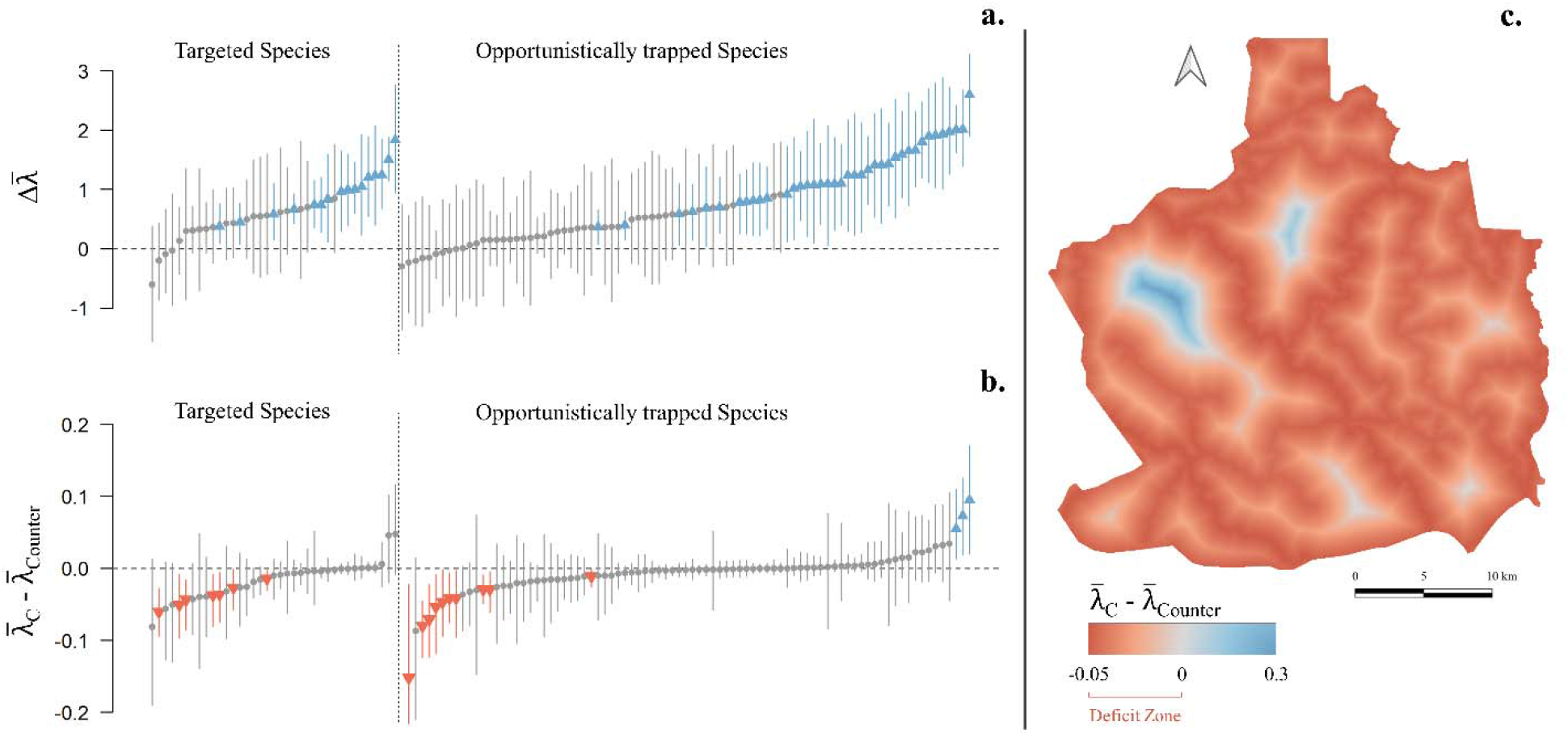
The recovery of bird species abundance in Harapan and the impacts it sustained from the trapping pressure. (a) Changes in mean species abundance between the early and current periods 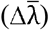. (b) The difference between mean species abundance in the current period and the counterfactual (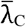 and 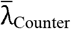); negative values indicate a deficit in abundance recovery attributed to the intensification of trapping. (c) Spatial patterns of the community-level recovery deficit 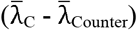 predicted in relation to the current access difficulty at Harapan. Deficit zone (red) are areas within Harapan where 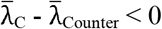 indicating areas where species’ abundance recovery linked to improvements in forest conditions could not compensate for the decline caused by trapping. For panels (a) and (b) each bar represents Bayesian equal-tailed credible intervals (89% ETI) of one species, belonging to either a guild targeted by trade: targeted or opportunistically trapped species. Species with significantly positive values (89% ETI > 0) are in blue (median values shown in triangles pointing up), species with significantly negative values (89% ETI < 0) are in red (median values shown in triangles pointing down), and species with non-significant values are in grey (median values shown in grey circle). Species are ordered by their increasing values within each guild.

Coinciding with increased human accessibility, we found that the negative impacts of trapping on bird species abundance had most likely intensified over time. First, the proportion of species whose abundance significantly increased with increasing access difficulty (i.e., 89% ETI for β_2_ > 0) had tripled over the study period, increasing from 5.4% in the early period to 16.2% in the current period (Fig. 3). In comparison, the proportion of species whose abundance significantly decreased with increasing access difficulty (i.e., 89% ETI for β_2_ < 0) declined from 6.9% to 1.5% between the two periods (Fig. 3). Second, 48.4% of species increased in the degree of association of their abundance with increasing access difficulty, by either changing from not significantly positive in the early period to significantly positive in the current period, or by becoming more positive in its median and narrower in 89% ETI over time (Fig. 3). At the community level, β_2_ increased slightly over time between early (median: -0.04 with 89% ETI: - 0.47 to 0.37) and current periods (median: 0.25 with 89% ETI: -0.68 to 1.17).

**Figure 3:**
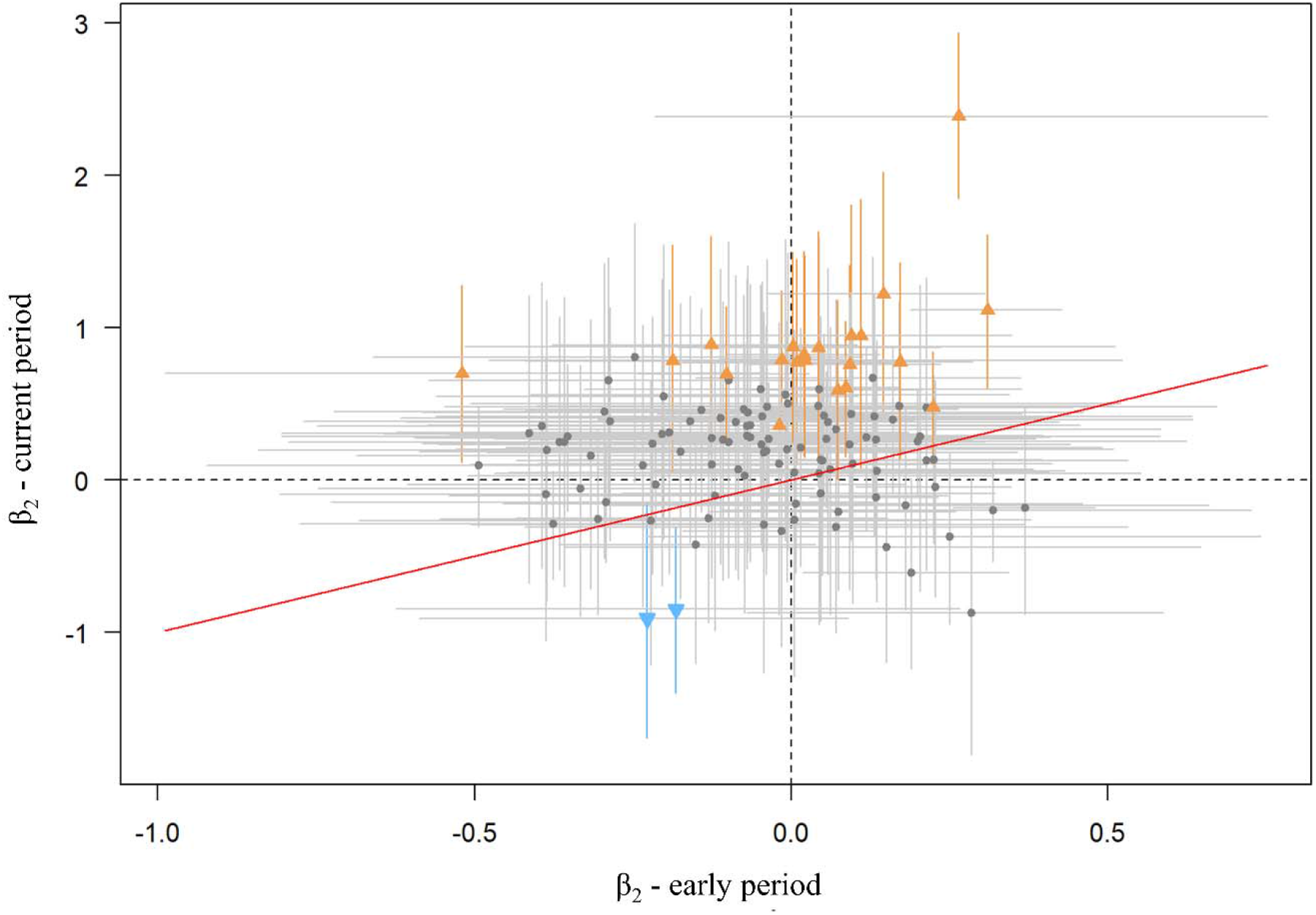
Estimated relationship between species abundance and access difficulty (β_2_). Bars represent 89% Bayesian equal-tailed credible intervals (89% ETI) of β_2_ for each species in in the early period (x-axis) and current period (y-axis). Species with significantly positive β_2_ (89% ETI for β_2_ > 0) in the current period are in orange (median values shown in triangles pointing up), species with significantly negative β_2_ in the current period (89% ETI for β_2_ < 0) are in blue (median values shown in triangles pointing down), and species with non-significant β_2_ are in grey (median values shown in grey circle). Red line represents 1:1 line, and the gray dashed lines represent zero response. A positive β_2_ indicates higher abundance in less accessible areas, likely linked to the negative impacts of trapping. β_2_ becoming more positive over time (median values above the red line), indicates intensified negative impacts of trapping on species abundance.

We found that 15% of the species analyzed exhibited a significant recovery deficit (i.e., negative difference between 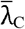 and 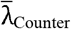, with 89% ETI < 0; Fig. 2b). In contrast, 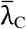 was greater than 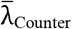 for only 2.3% of the species analyzed. This deficit was primarily driven by reductions in bird abundances in areas of greater human accessibility across Harapan, as demonstrated by the concentration of the ‘deficit zone’ of avian abundance recovery (i.e., region where 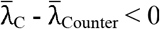) within ∼1 km from easy-access points at the community level (mean width of the deficit zone from access points: 475 m, range width: 0 – 1117 m; Fig. 2c).

A similar proportion of targeted and opportunistically trapped species exhibited positive changes in mean abundance between study periods 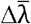; Table 1). Relative to opportunistically trapped species, a higher proportion of targeted species exhibited significantly positive β_2_ in the current period and a significant recovery deficit as represented by negative difference between 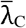 and 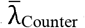 (Table 1). We found no difference in the width of the deficit zones for targeted (mean width of the deficit zone from access points: 473 m, range width: 0 – 1093 m) versus opportunistically trapped species (mean width of the deficit zone from access points: 477 m, range width: 0 – 1136 m). We found that a similar proportion of forest-dependent and generalist species exhibited positive changes in mean abundance 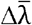 between study periods (Table 1). Relative to generalist species, we found a higher proportion of forest-dependent species exhibited significantly positive β_2_ in the current period and greater recovery deficits (Table 1).

**Table 1.**
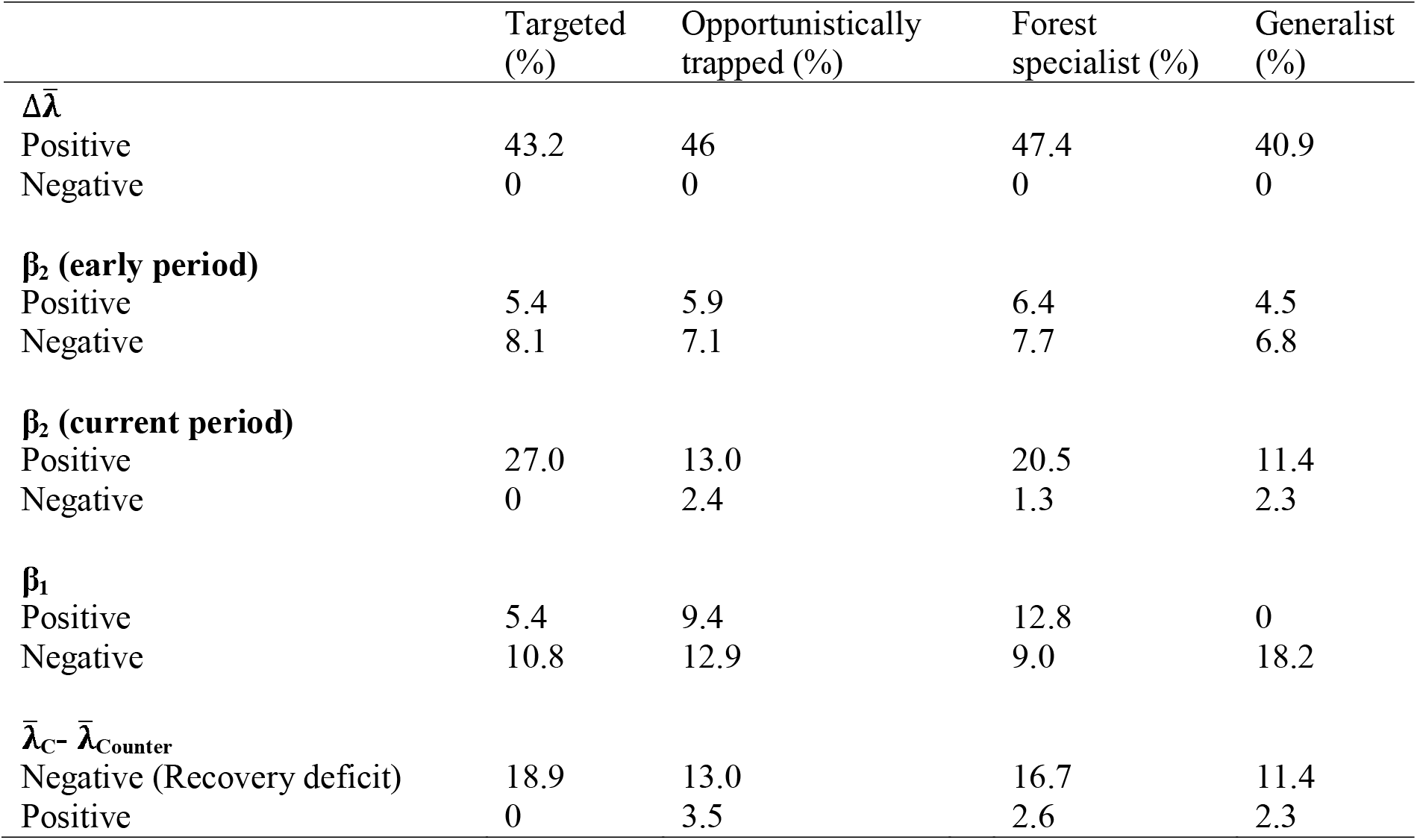
Proportion of species showing a significant effect. The effect is positive if the 89% Bayesian equal-tailed credible interval of the model derived posterior distributions are above 0 (i.e., 89% ETI > 0) and negative if 89% ETI < 0. 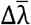 refers to changes in mean species abundance between the early and current periods, where a positive change shows recovery in bird species abundance. β_2_ is model-estimated relationship between species abundance and access difficulty, where positive β_2_ indicates higher abundance in less accessible areas, likely linked to the negative impacts of trapping. β_1_ is model-estimated relationship between species abundance and top-of-canopy height, where positive β_2_ indicates higher abundance in areas with tall canopy, likely linked to forest recovery. 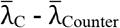 refers to changes in mean species abundance between current and counterfactual current period respectively, where negative values of 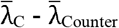 indicate a deficit in its abundance recovery (recovery deficit) attributed to the intensification of threats posed by trapping over time.

Finally, comparing indications of the negative impacts of trapping on species abundance as obtained from our field data against current IUCN threat assessments, we found that for the 15 species negatively affected by trapping in Harapan, exploitation or wildlife trade has yet to be formally recognized as a conservation threat by the IUCN Red List (Table S4). For the 21 species that exhibited either significant recovery deficit (i.e., 89% ETI for 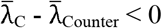) or whose abundance significantly increased with increasing access (i.e., 89% ETI for β_2_ < 0), most of them are currently classed as Least Concern (52% of species) or Near Threatened (33% of species; Table S4), and trapping is formally recognized by IUCN as a threat for only 6 of them.

## 4. DISCUSSION AND CONCLUSION

Our study shows that bird abundance significantly increased during a decade of forest regeneration and protection at a flagship site of tropical forest restoration in the lowlands of Sumatra. However, we also found indications of increasingly strong impacts of bird trapping on species abundance, which could hamper the conservation success of forest restoration. While most species showed signs of recovery over almost ten years of forest restoration at Harapan (Fig. 2a), for at least 16.2% of species, the intensifying trapping pressure (Fig. 3) dampened this recovery (Fig. 2b & c). Species prized in the pet trade and those dependent on late-successional forest habitat were particularly concentrated in remote, inaccessible areas (Table 1). We also found that although trapping is likely to significantly impact bird species in Indonesia, the current IUCN Red List (IUCN, 2019) considers trapping a threat to only a few of them (Table S4). Our findings demonstrate the potential of forest protection and regeneration in recovering Southeast Asia’s avian diversity but highlight the urgency of tackling the intensifying threat of pet trade.

Given appropriate conditions and recovery time, tropical forest restoration can allow biodiversity to recover, potentially to levels close to those found in primary forests (Crouzeilles et al., 2016; Gilroy et al., 2014; Rozendaal et al., 2019). Our results suggest that avian population recovery could happen rapidly within regenerating forests in lowland Sumatra, with notable abundance increases observed within a 10-year period. However, significant conservation gains are only possible if the recovering biodiversity is not jeopardized by factors beyond the scope of restoration. In the case of Harapan, financial and personnel resources supporting conservation have totaled approximately 20 million USD, with an annual operating cost of 1.48-2.5 million USD (Buergin, 2016; Diana and Jong, 2018; Silalahi et al., 2017), similar to the Indonesian average of 1 million USD required to manage per 40,000 Ha of forest restoration concession (Harrison et al., 2020). It is encouraging that this investment is associated with broad recovery in forest conditions and overall avian populations (Fig. 2a), but the level of recovery is likely to be lower than could have been achieved in the absence of growing levels of trapping (Benítez-López et al., 2017; Morton et al., 2021). As many bird species play key roles in the regeneration and functioning of forest ecosystems (e.g. through seed dispersal, Morrison & Lindell 2012; de la Peña-Domene et al. 2014), declines driven by trapping could lead to cascading ecological impacts that further limit the effectiveness of forest restoration (Gardner et al., 2019).

Our study adds to the growing evidence that increasing human incursion into forests, in particular for trade-driven trapping, constitutes a significant threat to Indonesia’s forest avifauna (Symes et al., 2018). Evidence from market surveys and communications with local experts, Harapan field staff and the trappers themselves, suggest that bird trapping is generally indiscriminate in its methods, in part because markets accept most species and because there is high mortality of birds along the supply chain (Chng et al., 2018b, 2015; Shepherd, 2006; Shepherd et al., 2004). These factors together incentivize trappers to maximize capture rates (Jepson and Ladle, 2005; Shepherd et al., 2004). As trapping depletes local bird populations, the economic incentive for indiscriminately trapping any species will likely intensify, with or without increases in market demand (Beastall et al., 2016; Courchamp et al., 2006; Shepherd, 2012; TRAFFIC, 2018). This could possibly explain the proportion of opportunistically trapped species showing negative effects of trapping pressure in our dataset (Table 1). The impacts of trapping on wildlife populations are further exacerbated by the rapid loss of forest habitat across Southeast Asia, which not only directly threatens biodiversity persistence, but also facilitates human access by expanding the road and trail network (Harris et al., 2017; Hughes, 2018; Margono et al., 2012).

Multiple threatened species detected in our surveys showed significant negative impacts of trapping pressure, and we found several species for which trapping has yet to be formally recognized as a conservation threat (Table S3). As an example, aside from species such as the Endangered Greater Green Leafbird, *Chloropsis sonnerati*, for which the threat of trapping has been well recognized (Eaton et al., 2017), our models suggested that the Dark-necked Tailorbird, *Orthotomus atrogularis*, Blue-crowned Hanging-Parrot, *Loriculus galgulus*, and Asian Fairy-bluebird, *Irena puella* (all Least Concern), may be also negatively affected by trapping (Table S3). In addition, a large number of Least Concern and Near Threatened species exhibited significant negative signals of trapping pressure (Table S4), possibly indicating looming population declines if the current trend continues.

Three caveats to our findings warrant discussion. First, because our early and current bird surveys were conducted by different surveyors, and at different survey time lengths, the observed difference in species abundance between study periods may have been influenced by these effects. We have taken measures to statistically alleviate this potential issue to the extent possible. Second, while we were able to assess the increases in TCH for low forest canopies (typically < 15 m) accurately, the potential TCH increases of tall forests were more challenging to assess due to the known issue of saturation of tree height predictions using LiDAR data (Hansen et al., 2016; Swinfield et al., 2019). Additionally, the resampled point count locations were from forested portions that did not experience fire and deforestation. This may be at least in part responsible for the observed weak relationship between species abundance and mean TCH. Third, we did not model the resulting landscape configuration from subsequent forest recovery and degradation, which may have influenced the estimation of bird population recovery (Bhakti et al., 2018; Carrara et al., 2015; Morante-Filho et al., 2021).

The threats that trapping poses to forest birds call for urgent conservation intervention. Here we provide several recommendations for Indonesia. First, the incentive to trap should be decreased by effective anti-trapping/poaching patrols, law enforcement of illegal selling of birds in markets, and including penalties for lawbreakers as deterrents that is coupled with alternative livelihood schemes (Leupen et al., 2018; López-Bao et al., 2015; Miller et al., 2019). Employing local communities and where possible, bird trappers in patrolling and as birdwatching guides could provide them with economic incentives to forego trapping and/or engage in conservation, and to cultivate a genuine long-term interest in supporting conservation (Widmann and Widmann, 2008). Second, region-wide threat assessments of the severity and extent of trade and trapping should be conducted on all bird species in Southeast Asia and check if localized threats we detected reflect the broader situation. Such assessments should use a combination of market- and field-based surveys to inform site-specific, targeted conservation interventions, such as *in situ* management of species, habitat (including nest site provisioning) and conservation breeding (Collar et al., 2012; M. Eaton et al., 2015; Harris et al., 2015; Kurniandaru, 2008; Pain et al., 2006). These assessments should also pre-emptively cover species not yet found in large quantities in markets and should consider potential taxonomic changes (Eaton et al., 2016). Third, behavioral change interventions that target consumers and other actors in the trade supply chain should be conducted, for example by encouraging competition categories exclusively for birds from commercial captive breeding or ‘ranching’, particularly for high-profile species such as the White-rumped Shama, *Copsychus malabaricus* (Burivalova et al., 2017; Jepson et al., 2017; Veríssimo, 2013). Efforts should be made to better understand and address the underlying drivers of wild bird trapping through a collaborative, multi-stakeholder approach, such as that showcased by the Asian Songbird Trade Specialist Group (Burivalova et al., 2017; Marshall et al., 2019; Shepherd and Cassey, 2017).

Forest restoration is urgently needed in many tropical regions that have experienced extensive deforestation and forest degradation, including Southeast Asia (Edwards et al., 2019; FAO and UNEP, 2020; Wilcove et al., 2013). However, its effectiveness in terms of biodiversity recovery could be compromised by wildlife trapping and exploitation. The increased accessibility of degraded forests compounds this challenge (Hughes, 2018). The realization of the conservation promise of forest restoration therefore hinges critically not only on effective restoration actions but also on addressing wildlife trapping.

## ACKNOWLEDGEMENTS

We gratefully acknowledge the funding support from the joint AEC-OBC Conservation Grant (No. P1250) provided by the Oriental Bird Club (OBC) and the Ecology Arboriculture Landscape’s (AEC), Sir Philip Reckitt Educational Trust (SPRET) travel grant, and European Commission’s Joint Master’s Degree Fellowship (FPA 2023 -- 0224/ 532524-1-FR-2012-1-ERA MUNDUS-EMMC). We sincerely thank S. Kumaran, F. Syamsuri, F. Hasudungan, and PT Restorasi Ekosistem Indonesia (PT REKI) for helping us to acquire research permits (126/SIP/FRP/E5/Dit.KVIV/2018) and for providing logistical support throughout the fieldwork. We thank Iwan and Andi for their assistance while collecting bird community data in the field. We also thank the anonymous reviewers for their constructive comments. H.S.S.C. thanks M. Persche and B.R. Shrestha for their detailed and helpful comments on the manuscript. H.S.S.C. thanks M. Persche for support while working on finishing the manuscript.

## REFERENCES

Asner GP, Brodrick PG, Philipson C, Vaughn NR, Martin RE, Knapp DE, Heckler J, Evans LJ, Jucker T, Goossens B. 2018. Mapped aboveground carbon stocks to advance forest conservation and recovery in Malaysian Borneo. Biological Conservation 217:289–310. DOI: 10.1016/j.biocon.2017.10.020.

Barlow, J., França, F., Gardner, T.A., Hicks, C.C., Lennox, G.D., Berenguer, E., Castello, L., Economo, E.P., Ferreira, J., Guénard, B., Gontijo Leal, C., Isaac, V., Lees, A.C., Parr, C.L., Wilson, S.K., Young, P.J., Graham, N.A.J., 2018. The future of hyperdiverse tropical ecosystems. Nature 559, 517–526. DOI: 10.1038/s41586-018-0301-1

Beastall C, Shepherd CR, Hadiprakarsa Y, Martyr D. 2016. Trade in the Helmeted Hornbill Rhinoplax vigil: the ‘ivory hornbill.’ Bird Conservation International 26:137–146. DOI: 10.1017/S0959270916000010.

Benítez-López, A., Alkemade, R., Schipper, A.M., Ingram, D.J., Verweij, P.A., Eikelboom, J.A.J., Huijbregts, M.A.J., 2017. The impact of hunting on tropical mammal and bird populations. Science. 356, 180–183. DOI: 10.1126/science.aaj1891

Beygelzimer A, Kakadet S, Langford J, Arya S, Mount D, Li S. 2019. Package ‘FNN’: FNN: Fast Nearest Neighbor Search Algorithms and Applications. Version 1.1.3. Available from https://cran.r-project.org/ (accessed October 2020).

Bhakti, T., Goulart, F., de Azevedo, C.S., Antonini, Y. 2018. Does scale matter? The influence of three-level spatial scales on forest bird occurrence in a tropical landscape. PLoS One 13. DOI: 10.1371/journal.pone.0198732.

Billerman SM, Keeney BK, Rodewald PG, Schulenberg TS, editors. 2020. Birds of the World. Cornell Laboratory of Ornithology, Ithaca, NY. Available from https://birdsoftheworld.org/ (accessed November 2020).

BirdLife International. 2020. Species factsheet: Rhinoplax vigil. Available from http://www.birdlife.org/ (accessed November 2020).

BirdLife International. 2017. Forests of Hope site - Harapan Rainforest, Indonesia. Available from http://www.birdlife.org/ (accessed November 2020).

Breiman L, Cutler A. 2018. Package ‘randomForest’: Breiman and Cutler’s Random Forests for Classification and Regression. Version 4.6. Available from https://cran.r-project.org/ (accessed October 2019).

Buergin R. 2016. Ecosystem restoration concessions in Indonesia: conflicts and discourses. Critical Asian Studies 48:278–301. DOI: 10.1080/14672715.2016.1164017.

Burivalova Z, Lee TM, Hua F, Lee JSH, Prawiradilaga DM, Wilcove DS. 2017. Understanding consumer preferences and demography in order to reduce the domestic trade in wild-caught birds. Biological Conservation 209:423–431. DOI: 10.1016/j.biocon.2017.03.005.

Carrara, E., Arroyo-Rodríguez, V., Vega-Rivera, J.H., Schondube, J.E., de Freitas, S.M., Fahrig, L. 2015. Impact of landscape composition and configuration on forest specialist and generalist bird species in the fragmented Lacandona rainforest, Mexico. Biological Conservation 184: 117–126. DOI: 10.1016/j.biocon.2015.01.014

Chazdon, R., Brancalion, P. 2019. Restoring forests as a means to many ends. Science. 365: 24–25. DOI: 10.1126/science.aax9539.

Chazdon RL, Brancalion PHS, Lamb D, Laestadius L, Calmon M, Kumar C. 2017. A Policy-Driven Knowledge Agenda for Global Forest and Landscape Restoration. Conservation Letters 10:125–132. DOI: 10.1111/conl.12220.

Chng SCL, Eaton JA, Krishnasamy K, Shepherd CR. 2015. In the Market for Extinction: An inventory of Jakarta’s bird markets. TRAFFIC Report. Petaling Jaya, Selangor, Malaysia.

Chng, S.C.L., Guciano, M., Eaton, J.A., 2016. In the market for extinction: Sukahaji, Bandung, Java, Indonesia. BirdingASIA 26: 22–28.

Chng, S.C.L., Krishnasamy, K., Eaton, J.A., 2018a. In the market for extinction: the cage bird trade in Bali. Forktail 34: 35–41.

Chng, S.C.L., Shepherd, C.R., Eaton, J.A., 2018b. In the market for extinction: birds for sale at selected outlets in Sumatra. TRAFFIC Bulletin 30(1):15–22.

Collar NJ, Gardner L, Jeggo DF, Marcordes B, Owen A, Pagel T, Pes T, Vaidl A, Wilkinson R, Wirth R. 2012. Conservation breeding and the most threatened birds in Asia. BirdingAsia 18:50–57.

Cosset CCP, Edwards DP. 2017. The effects of restoring logged tropical forests on avian phylogenetic and functional diversity. Ecological Applications 27:1932–1945. DOI: 10.1002/eap.1578.

Courchamp F, Angulo E, Rivalan P, Hall RJ, Signoret L, Bull L, Meinard Y. 2006. Rarity value and species extinction: the anthropogenic Allee effect. PLOS Biology 4. DOI: 10.1371/journal.pbio.0040415.

Crouzeilles R, Curran M, Ferreira MS, Lindenmayer DB, Grelle CEV, Rey Benayas JM. 2016. A global meta-Analysis on the ecological drivers of forest restoration success. Nature Communications 7. DOI: 10.1038/ncomms11666

Csillik O, Kumar P, Mascaro J, O’Shea T, Asner GP. 2019. Monitoring tropical forest carbon stocks and emissions using Planet satellite data. Scientific Reports 9:17831. DOI: 10.1038/s41598-019-54386-6

de la Peña-Domene M, Martínez-Garza C, Palmas-Pérez S, Rivas-Alonso E, Howe HF. 2014. Roles of birds and bats in early tropical-forest restoration. PloS one 9: e104656. DOI: 10.1371/journal.pone.0104656.

Diana E, Jong HN. 2018. End of funding dims hopes for a Sumatran forest targeted by palm oil growers. Mongabay. Available from https://news.mongabay.com/ (accessed March 2020).

Eaton JA, Chng SCL, Miller AE. 2017. Second South-east Asian Songbird Crisis Summit. TRAFFIC Bulletin 29(1):3–4.

Eaton, J.A., Shepherd, C.R., Rheindt, F.E., Harris, J.B.C., van Balen, S., Wilcove, D.S., Collar, N.J. 2015. Trade-driven extinctions and near-extinctions of avian taxa in Sundaic Indonesia. Forktail: 1–12.

Eaton JA, van Balen S, Brickle NW, Rheindt FE. 2016. Birds of the Indonesian Archipelago: Greater Sundas and Wallacea. Lynx. Spain.

Eaton, M., Aebischer, N., Brown, A., Hearn, R., Lock, L., Musgrove, A., Noble, D., Stroud, D., Gregory, R. 2015. Birds of Conservation Concern 4: the population status of birds in the UK, Channel Islands and Isle of Man. British Birds 108: 708–746.

Edwards DP, Ansell FA, Ahmad AH, Nilus R, Hamer KC. 2009. The value of rehabilitating logged rainforest for birds. Conservation Biology 23:1628–1633. DOI: 10.1111/j.1523-1739.2009.01330.x

Edwards, D.P., Magrach, A., Woodcock, P., Ji, Y., Lim, N.T.L., Edwards, F.A., Larsen, T.H., Hsu, W.W., Benedick, S., Khen, C.V., Chung, A.Y.C., Reynolds, G., Fisher, B., Laurance, W.F., Wilcove, D.S., Hamer, K.C., Yu, D.W. 2014. Selective-logging and oil palm: Multitaxon impacts, biodiversity indicators, and trade-offs for conservation planning. Ecological Applications 24:2029–2049. DOI: 10.1890/14-0010.1

Edwards DP, Socolar JB, Mills SC, Burivalova Z, Koh LP, Wilcove DS. 2019. Conservation of Tropical Forests in the Anthropocene. Current Biology 29: R1008–R1020.

FAO, UNEP. 2020. The State of the World’s Forests 2020. Forests, biodiversity and people. Accessed from http://www.fao.org/ (accessed August 2020).

Farnsworth GL, Pollock KH, Nichols JD, Simons TR, Hines JE, Sauer JR. 2002. A removal model for estimating detection probabilities from point-count surveys. The Auk 119:414–425.

Gardner CJ, Bicknell JE, Baldwin-Cantello W, Struebig MJ, Davies ZG. 2019. Quantifying the impacts of defaunation on natural forest regeneration in a global meta-analysis. Nature Communications 10:4590. DOI: 10.1038/s41467-019-12539-1.

Gibson, L., Lee, T.M., Koh, L.P., Brook, B.W., Gardner, T.A., Barlow, J., Peres, C.A., Bradshaw, C.J.A., Laurance, W.F., Lovejoy, T.E., Sodhi, N.S. 2011. Primary forests are irreplaceable for sustaining tropical biodiversity. Nature 478, 378–381. DOI: 10.1038/nature10425.

Gilroy JJ, Woodcock P, Edwards FA, Wheeler C, Baptiste BLG, Uribe CAM, Haugaasen T, Edwards DP. 2014. Cheap carbon and biodiversity co-benefits from forest regeneration in a hotspot of endemism. Nature Climate Change 4:503–507. DOI: 10.1038/nclimate2200.

Grant PR, Grant BR, Huey RB, Johnson MTJ, Knoll AH, Schmitt J. 2017. Evolution caused by extreme events. Philosophical Transactions of the Royal Society B: Biological Sciences 372:20160146.

Hansen MC, Potapov P V, Goetz SJ, Turubanova S, Tyukavina A, Krylov A, Kommareddy A, Egorov A. 2016. Mapping tree height distributions in Sub-Saharan Africa using Landsat 7 and 8 data. Remote Sensing of Environment 185:221–232. DOI: 10.1016/j.rse.2016.02.023.

Harris JBC, Green JMH, Prawiradilaga DM, Giam X, Hikmatullah D, Putra CA, Wilcove DS. 2015. Using market data and expert opinion to identify overexploited species in the wild bird trade. Biological Conservation 187:51–60. DOI: 10.1016/j.biocon.2015.04.009.

Harris, J.B.C., Tingley, M.W., Hua, F., Yong, D.L., Adeney, J.M., Lee, T.M., Marthy, W., Prawiradilaga, D.M., Sekercioglu, C.H., Suyadi, C.H., Winarni, N., Wilcove, D.S. 2017. Measuring the impact of the pet trade on Indonesian birds. Conservation Biology 31:394–405. DOI: 10.1111/cobi.12729.

Harrison RD, Swinfield T. 2015. Restoration of Logged Humid Tropical Forests: An Experimental Programme at Harapan Rainforest, Indonesia. Tropical Conservation Science 8:4–16. DOI: 10.1177/194008291500800103.

Harrison, R.D., Swinfield, T., Ayat, A., Dewi, S., Silalahi, M., Heriansyah, I., 2020. Restoration concessions: a second lease on life for beleaguered tropical forests? Frontiers in Ecology and the Environment 18: 567–575. DOI: 10.1002/fee.2265.

Hastie T, Tibshirani R, Friedman J. 2009. The elements of statistical learning: data mining, inference, and prediction. Springer Science & Business Media. Springer-Verlag, New York.

Hua F, Yong DL, Janra MN, Fitri LM, Prawiradilaga D, Sieving KE. 2016. Functional traits determine heterospecific use of risk-related social information in forest birds of tropical South-East Asia. Ecology and Evolution 6:8485–8494. DOI: 10.1002/ece3.2545.

Hughes, A.C. 2021. Wildlife trade. Current Biology 31: 1218–1224. DOI: 10.1016/j.cub.2021.08.056

Hughes AC. 2018. Have Indo-Malaysian forests reached the end of the road? Biological Conservation 223:129–137. DOI: 10.1016/j.biocon.2018.04.029.

IUCN. 2019. The IUCN Red List of Threatened Species. Available from http://www.iucnredlist.org (accessed October 2020).

Jepson P. 2010. Towards an Indonesian bird conservation ethos: reflections from a study of bird-keeping in the cities of Java and Bali. Pages 313–330 in Tidemann SC, Gosler A, editors. Ethno-ornithology: Birds, indigenous peoples, culture and society: Routledge, UK.

Jepson, P., Ladle, R.J. 2005. Bird-keeping in Indonesia: Conservation impacts and the potential for substitution-based conservation responses. Oryx 39: 442–448. DOI: 10.1017/S0030605305001110

Jin S, Sader SA. 2005. Comparison of time series tasseled cap wetness and the normalized difference moisture index in detecting forest disturbances. Remote sensing of Environment 94:364–372. DOI: 10.1016/j.rse.2004.10.012.

Kruschke, J. 2014. Doing Bayesian data analysis: A tutorial with R, JAGS, and Stan.

Kurniandaru S. 2008. Providing nest boxes for Java sparrows Padda oryzivora in response to nest site loss due to building restoration and an earthquake, Prambanan Temple, Java, Indonesia. Conservation Evidence 5:62–68.

Latja P, Valtonen A, Malinga GM, Roininen H. 2016. Active restoration facilitates bird community recovery in an Afrotropical rainforest. Biological Conservation 200:70–79. DOI: 10.1016/j.biocon.2016.05.035.

Lee DC, Lindsell JA. 2011. Biodiversity of Harapan Rainforest: Summary report on baseline surveys of mammals, birds, fish, herptiles, butterflies and habitat. Royal Society for Protection of Birds. Sandy, UK.

Lee, D.C., Powell, V.J., Lindsell, J.A. 2019. Understanding landscape and plot-scale habitat utilisation by Malayan sun bear (Helarctos malayanus) in degraded lowland forest. Acta Oecologica 96: 1–9. DOI: 10.1016/j.actao.2019.02.002

Lee, D.C., Powell, V.J., Lindsell, J.A., 2014. The conservation value of degraded forests for agile gibbons Hylobates agilis. American Journal of Primatology 77: 76–85. DOI: 10.1002/ajp.22312

Leupen BTC, Krishnasamy K, Shepherd CR, Chng SCL, Bergin D, Eaton JA, Yukin DA, Hue SKP, Miller A, Nekaris KA-I. 2018. Trade in White-rumped Shamas Kittacincla malabarica demands strong national and international responses. Forktail Journal of Asian Ornithology 34:1–8.

Lewis SL, Wheeler CE, Mitchard ETA, Koch A. 2019. Restoring natural forests is the best way to remove atmospheric carbon. Nature 568:25–28. DOI: 10.1038/d41586-019-01026-8

López-Bao, J. V, Blanco, J.C., Rodríguez, A., Godinho, R., Sazatornil, V., Álvares, F., Garcia, E., Llaneza, L., Rico, M., Cortés, Y., Palacios, V., Chapron, G. 2015. Toothless wildlife protection laws. Biodiversity and Conservation 24:2105–2108. DOI: 10.1007/s10531-015-0914-8

Makowski, D., Ben-Shachar, M.S., Lüdecke, D. 2019. bayestestR: Describing Effects and their Uncertainty, Existence and Significance within the Bayesian Framework. Journal of Open Source Software 4(40),1541. DOI: 10.21105/joss.01541.

Margono BA, Turubanova S, Zhuravleva I, Potapov P, Tyukavina A, Baccini A, Goetz S, Hansen MC. 2012. Mapping and monitoring deforestation and forest degradation in Sumatra (Indonesia) using Landsat time series data sets from 1990 to 2010. Environmental Research Letters 7. DOI: 10.1088/1748-9326/7/3/034010.

Marshall H, Collar NJ, Lees AC, Moss A, Yuda P, Marsden SJ. 2019. Spatio-temporal dynamics of consumer demand driving the Asian Songbird Crisis. Biological Conservation:108237. DOI: 10.1016/j.biocon.2019.108237.

McElreath, R. 2018. Statistical Rethinking: A Bayesian Course with Examples in R and Stan.

Miller AE, Gary D, Ansyah J, Sagita N, Muflihati, Kartikawati, Adirahmanta SN. 2019. Socioeconomic Characteristics of Songbird Shop Owners in West Kalimantan, Indonesia. Tropical Conservation Science. DOI: 10.1177/1940082919889510.

Morante-Filho, J.C., Benchimol, M., Faria, D. 2021. Landscape composition is the strongest determinant of bird occupancy patterns in tropical forest patches. Landscape Ecology 36: 105–117. DOI: 10.1007/s10980-020-01121-6

Morrison EB, Lindell CA. 2012. Birds and bats reduce insect biomass and leaf damage in tropical forest restoration sites. Ecological Applications 22:1526–1534. DOI: 10.1890/11-1118.1.

Morton, O., Scheffers, B.R., Haugaasen, T., Edwards, D.P. 2021. Impacts of wildlife trade on terrestrial biodiversity. Nature Ecology and Evolution 5: 540–548. DOI: 10.1038/s41559-021-01399-y

Owen KC, Melin AD, Campos FA, Fedigan LM, Gillespie TW, Mennill DJ. 2020. Bioacoustic analyses reveal that bird communities recover with forest succession in tropical dry forests. Avian Conservation and Ecology 15 (1): 25. DOI: 10.5751/ACE-01615-150125.

Pain DJ, Martins TLF, Boussekey M, Diaz SH, Downs CT, Ekstrom JMM, Garnett S, Gilardi JD, McNiven D, Primot P. 2006. Impact of protection on nest take and nesting success of parrots in Africa, Asia and Australasia. Animal Conservation 9:322–330. DOI: 10.1111/j.1469-1795.2006.00040.x.

Plummer M. 2016. Package ‘rjags’: Bayesian graphical models using MCMC. Version 3. Available from https://cran.r-project.org/ (accessed November 2018).

Plummer M. 2012. JAGS Version 3.3.0 user manual. International Agency for Research on Cancer, Lyon, France.

Plummer M. 2003. JAGS: A program for analysis of Bayesian graphical models using Gibbs sampling. Proceedings of the 3rd international workshop on distributed statistical computing. Vienna, Austria.

Potapov, P., Hansen, M.C., Laestadius, L., Turubanova, S., Yaroshenko, A., Thies, C., Smith, W., Zhuravleva, I., Komarova, A., Minnemeyer, S., Esipova, E. 2017. The last frontiers of wilderness: Tracking loss of intact forest landscapes from 2000 to 2013. Science Advances 3. DOI: 10.1126/sciadv.1600821

R Development Core Team. 2018. R: a language and environment for statistical computing. R Foundation for Statistical Computing, Vienna.

Rentschlar, K.A., Miller, A.E., Lauck, K.S., Rodiansyah, M., Bobby, Muflihati, Kartikawati, 2018. A silent morning: The songbird trade in Kalimantan, Indonesia. Tropical Conservation Science. DOI: 10.1177/1940082917753909

Royle JA. 2004. N-Mixture Models for Estimating Population Size from Spatially Replicated Counts. Biometrics 60:108–115. DOI: 10.1111/j.0006-341X.2004.00142.x.

Rozendaal DMA, Bongers F, Aide TM, Alvarez-Dávila E, Ascarrunz N, Balvanera P, Becknell JM, Bentos T V, Brancalion PHS, Cabral GAL. 2019. Biodiversity recovery of Neotropical secondary forests. Science Advances 5. DOI: 10.1126/sciadv.aau3114.

Scheffers BR, Oliveira BF, Lamb I, Edwards DP. 2019. Global wildlife trade across the tree of life. Science 366:71–76. DOI: 10.1126/science.aav5327

Senior RA, Hill JK, Edwards DP. 2019. Global loss of climate connectivity in tropical forests. Nature Climate Change 9:623–626. DOI: 10.1038/s41558-019-0529-2.

Shepherd CR. 2012. The owl trade in Jakarta, Indonesia: a spot check on the largest bird markets. BirdingASIA 18:58–59.

Shepherd CR. 2006. The bird trade in Medan, north Sumatra: An overview. BirdingASIA 5:16–24.

Shepherd CR, Cassey P. 2017. Songbird trade crisis in Southeast Asia leads to the formation of IUCN SSC Asian Songbird Trade Specialist Group. Journal of Indonesian Natural History 5:3–5.

Shepherd, C.R., Eaton, J., Asia, B., Shepherd, C.R., Eaton, J.A., Chng, S.C.L. 2016. Pittas for a pittance: observations on the little known illegal trade in Pittidae in west Indonesia. BirdingASIA 24: 18–20.

Shepherd CR, Sukumaran J, Wich SA. 2004. Open season: An analysis of the pet trade in Medan, Sumatra, 1997-2001. TRAFFIC Southeast Asia Report. Petaling Jaya, Selangor, Malaysia.

Silalahi M, Utomo AB, Walsh TA, Ayat A, Bashir S. 2017. Indonesia’s ecosystem restoration concessions. Unasylva 68: 63–70.

Sodhi, N.S., Pin, L., Clements, R., Wanger, T.C., Hill, J.K., Hamer, K.C., Clough, Y., Tscharntke, T., Rose, M., Posa, C., Ming, T. 2010. Conserving Southeast Asian forest biodiversity in human-modified landscapes. Biological Conservation. DOI: 10.1016/j.biocon.2009.12.029

Strassburg, B.B.N., Iribarrem, A., Beyer, H.L., Cordeiro, C.L., Crouzeilles, R., Jakovac, C.C., Braga Junqueira, A., Lacerda, E., Latawiec, A.E., Balmford, A., Brooks, T.M., Butchart, S.H.M., Chazdon, R.L., Erb, K.-H., Brancalion, P., Buchanan, G., Cooper, D., Díaz, S., Donald, P.F., Kapos, V., Leclère, D., Miles, L., Obersteiner, M., Plutzar, C., de M. Scaramuzza, C.A., Scarano, F.R., Visconti, P. 2020. Global priority areas for ecosystem restoration. Nature. DOI: 10.1038/s41586-020-2784-9.

Su Y-S, Yajima M. 2015. Package ‘R2jags’: Using R to run ‘JAGS.’ Version 0.5-7. Available from https://cran.r-project.org/ (accessed November 2018).

Swinfield T, Lindsell JA, Williams J V, Harrison RD, Gemita E, Schönlieb CB, Coomes DA. 2019. Accurate Measurement of Tropical Forest Canopy Heights and Aboveground Carbon Using Structure from Motion. Remote Sensing 11:928. DOI: 10.3390/rs11080928

Symes WS, Edwards DP, Miettinen J, Rheindt FE, Carrasco LR. 2018. Combined impacts of deforestation and wildlife trade on tropical biodiversity are severely underestimated. Nature Communications 9:4052. DOI: 10.1038/s41467-018-06579-2.

TRAFFIC. 2018. Massive wild bird seizures reflect soaring pressure on Sumatran birds. Available from https://www.traffic.org/ (September 2019).

Veríssimo D. 2013. Influencing human behaviour: an underutilised tool for biodiversity management. Conservation Evidence 10:29–31.

Watson, J.E.M., Evans, T., Venter, O., Williams, B., Tulloch, A., Stewart, C., Thompson, I., Ray, J.C., Murray, K., Salazar, A., McAlpine, C., Potapov, P., Walston, J., Robinson, J.G., Painter, M., Wilkie, D., Filardi, C., Laurance, W.F., Houghton, R.A., Maxwell, S., Grantham, H., Samper, C., Wang, S., Laestadius, L., Runting, R.K., Silva-Chávez, G.A., Ervin, J., Lindenmayer, D. 2018. The exceptional value of intact forest ecosystems. Nature Ecology and Evolution 2: 599–610. DOI: 10.1038/s41559-018-0490-x

Widmann P, Widmann IL. 2008. The cockatoo and the community: ten years of Philippine Cockatoo Conservation Programme. Birding Asia 10:23–29.

Wilcove DS, Giam X, Edwards DP, Fisher B, Koh LP. 2013. Navjot’s nightmare revisited: logging, agriculture, and biodiversity in Southeast Asia. Trends in Ecology and Evolution 28:531–540. DOI: 10.1016/j.tree.2013.04.005

Xue J, Su B. 2017. Significant remote sensing vegetation indices: A review of developments and applications. Journal of Sensors. Hindawi. DOI: 10.1155/2017/1353691.

Yamaura Y, Andrew Royle J, Kuboi K, Tada T, Ikeno S, Makino S. 2011. Modelling community dynamics based on species□level abundance models from detection/non detection data. Journal of Applied Ecology 48:67–75.

Yamaura Y, Royle JA, Shimada N, Asanuma S, Sato T, Taki H, Makino S. 2012. Biodiversity of man-made open habitats in an underused country: a class of multispecies abundance models for count data. Biodiversity and Conservation 21:1365–1380. DOI: 10.1007/s10531-012-0244-z.

