## Supplementary material for "Avifauna recovers faster in areas less accessible to trapping in regenerating tropical forests": Supplemenatry Information

**Appendix A**: Quantifying forest condition as represented by TCH, using Landsat imagery and a model derived from the LiDAR dataset.

The LiDAR dataset was collected October 24, 2014 using a Leica ALS70 camera mounted on a Pilatus Porter aircraft flown at 800 m above ground at 100 km per hour. For the Landsat imagery, the five vegetation indices (VIs) we used were: (1) normalised difference vegetation index (NDVI); (2) global environmental monitoring index (GEMI); (3) soil-adjusted vegetation index (SAVI); (4) enhanced vegetation index (EVI); and (5) normalized difference moisture index (NDMI) (Jin & Sader 2005). We then accessed the entire Tier 1 Landsat Surface Reflectance catalog through Google Earth Engine (GEE), masked the low-quality pixels (e.g., clouds and shadows) using the cfmask pixel quality values (provided in GEE), calculated the VI values, and converted them into annual median values. For the SAVI index, we additionally described their spatial contrast by assessing horizontal variation in canopy structure, which we achieved by calculating grey-level covariance matrix values (mean and homogeneity) for 30 m and 90 m offsets using the package ‘glcm’ (version 1.6.5; Zvoleff, 2020) under program R (R Development Core Team 2018). When predicting TCH, we used two discrete Landsat images two years apart, which enabled us to sample the variation arising from differences in observation and vegetation conditions, which minimized over-fitting and provided an unbiased test of model accuracy when predicting to unseen Landsat images (Hastie et al. 2009). During the assessment of predicted TCH, 25% of the out-of-sample variance was explained with a root-mean-squared error of 5.27 m across the validation areal coverage of 2014.

**References:**

Hastie T, Tibshirani R, Friedman J. 2009. The elements of statistical learning: data mining, inference, and prediction. Springer Science & Business Media.

Jin S, Sader SA. 2005. Comparison of time series tasseled cap wetness and the normalized difference moisture index in detecting forest disturbances. Remote sensing of Environment **94**:364–372. Elsevier.

R Development Core Team. 2018. R: a language and environment for statistical computing. R Foundation for Statistical Computing, Vienna.

Zvoleff A. 2020. Package “glcm.” Available from https://cran.r-project.org/web/packages/glcm/glcm.pdf.

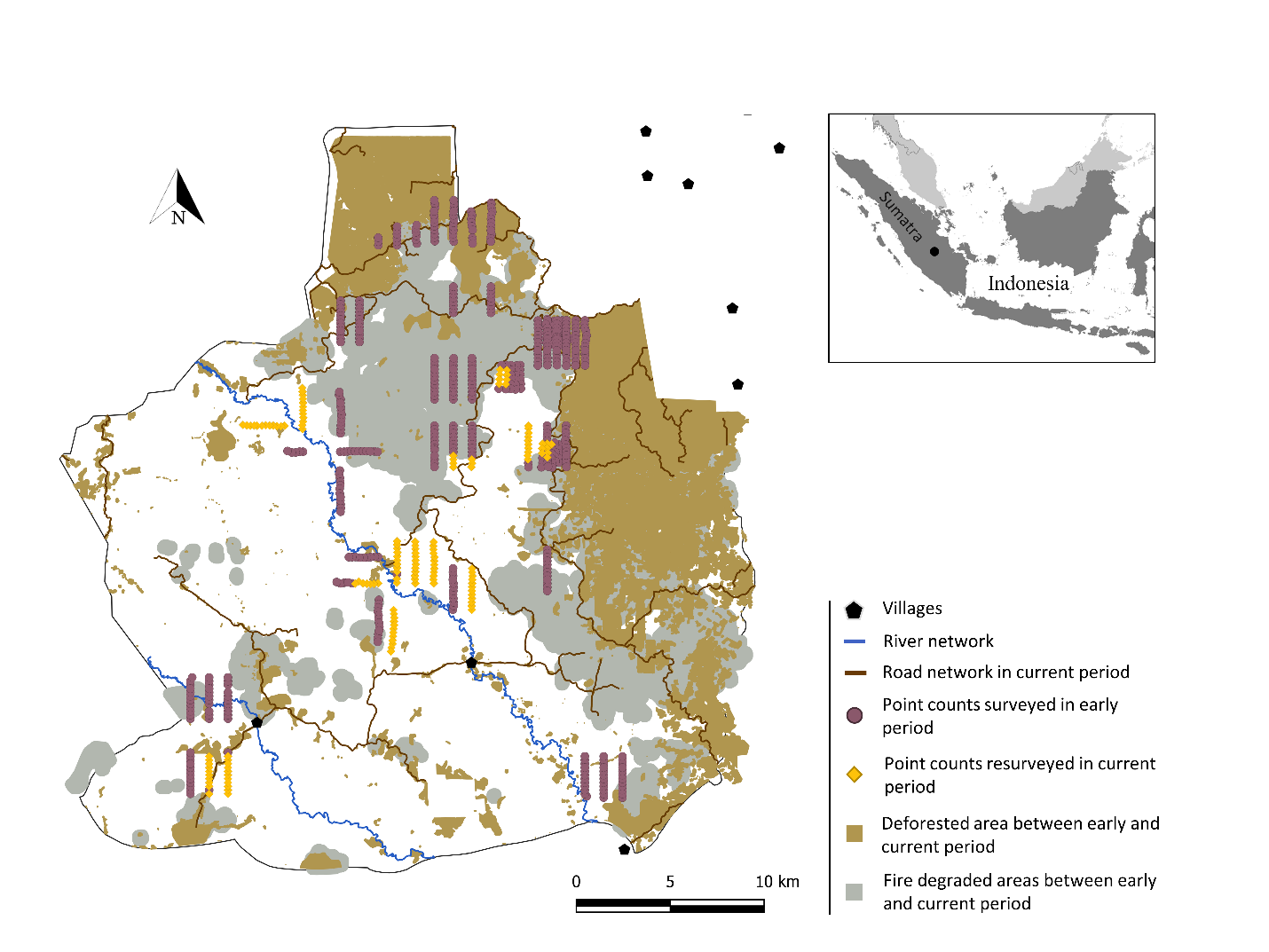

**Figure S1:** Map of anthropogenically degraded areas in Harapan between the early and current periods (2009-2011 and 2018, respectively), with deforestation (brown) and fire-induced degradation (grey). Point count stations were surveyed during the early period (purple) or in both the early and current periods (yellow).

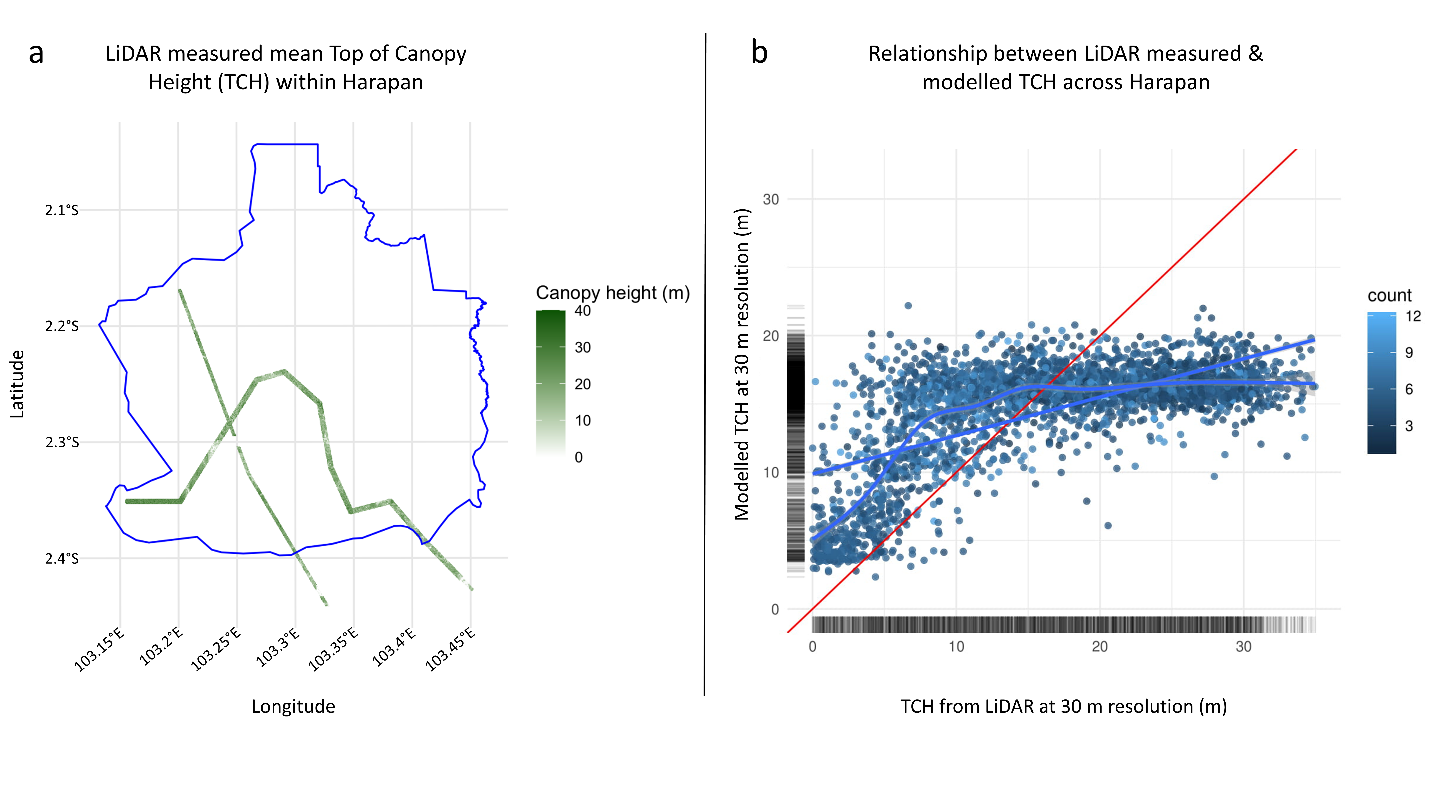

**Figure S2:** LiDAR measurement and Harapan-wide prediction of top-of-canopy height (TCH). (a) Map showing the flight path and the mean TCH (in meters) measured using LiDAR in 2014. (b) The out of set model validation for the random forest prediction of canopy height at 30 m resolution from Landsat imagery. The red line is the 1:1 line; the straight blue line is the linear fit between the predicted and observed; the curved blue line is the spline fit between the predicted and observed.

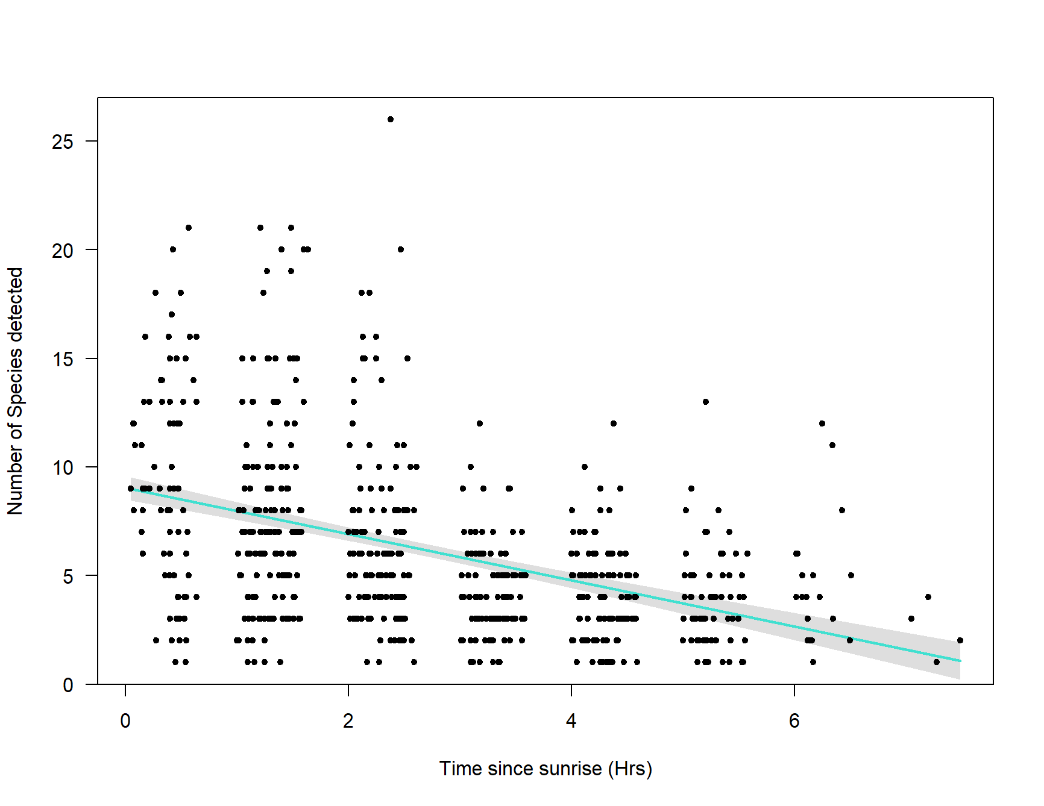

Number of Species Detected

**a.**

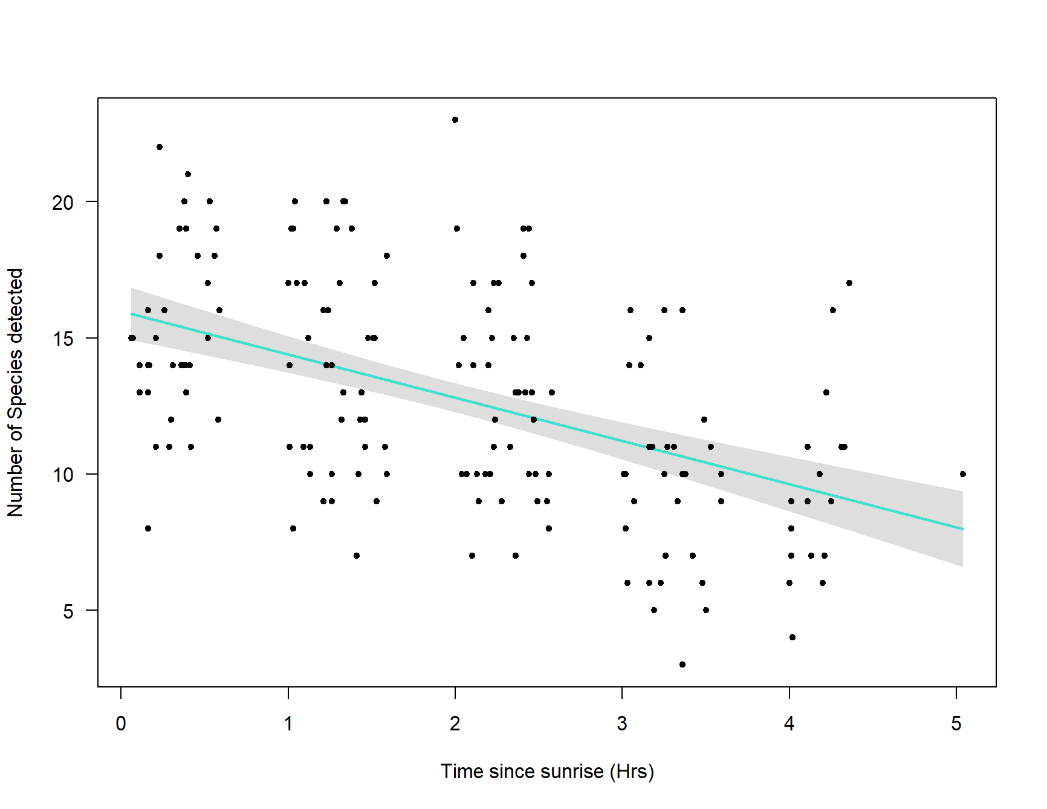

Number of Species Detected

Time since sunrise (Hrs)

**b.**

**Figure S3:** Relationship between Time since Sunrise (x-axis) and number of Species Detected at each point count location (y-axis) in, **a:** early period (y = -1.03x + 9.03, r^2^ = 0.18, p < 0.01) and **b:** current period (y = -1.58x + 15.97, r^2^ = 0.25, p < 0.01). Turquoise line represents the model fit and grey polygon represents the 95% confidence interval.

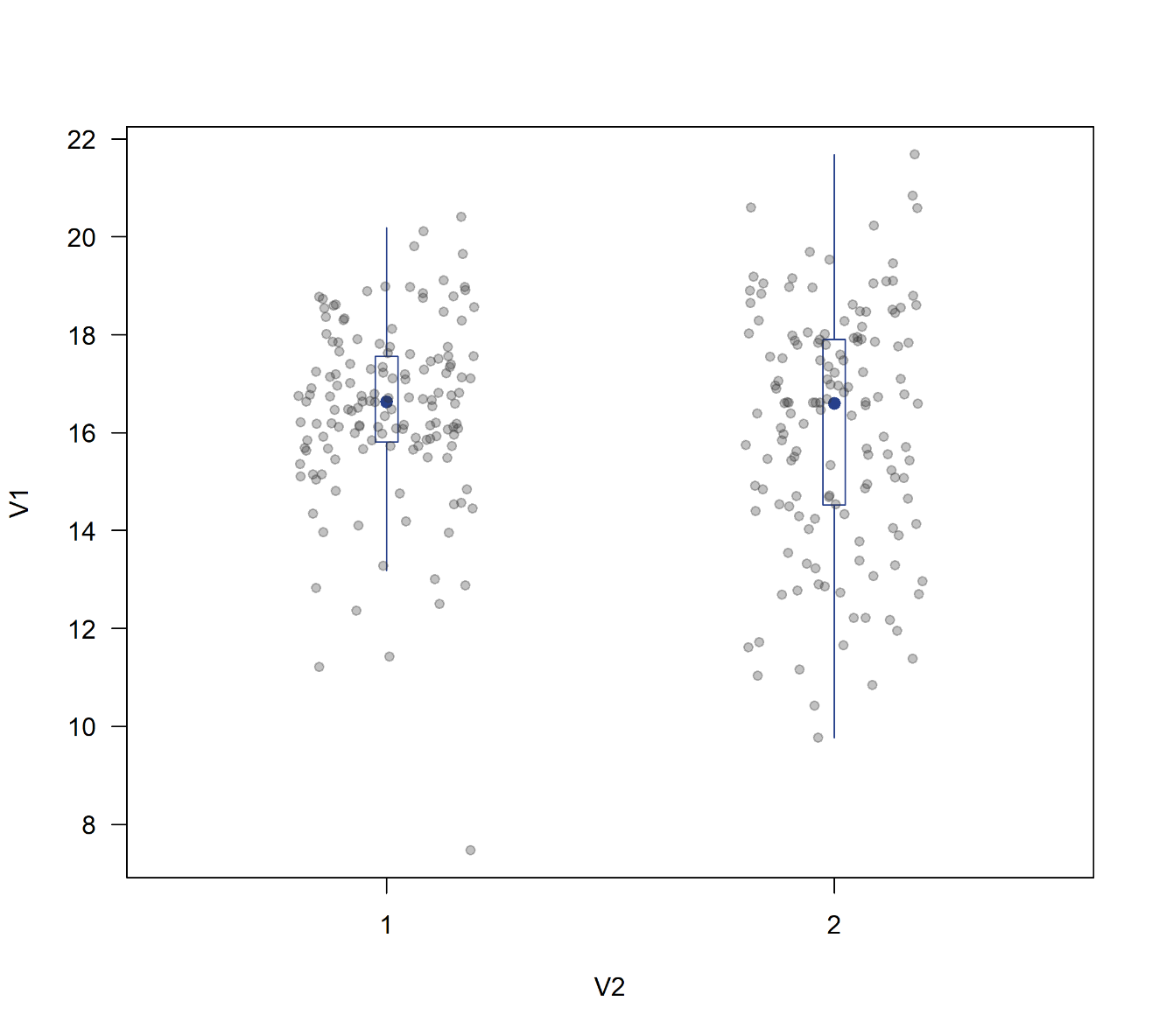

Top of Canopy Height (m)

Early Period

Current Period

**Figure S4**: Mean TCH across resampled locations between early and current periods. Mean difference = 0.38 with CI = -0.04, 0.79 (Paired t-test, t_20_ = 1.8, p = 0.08).

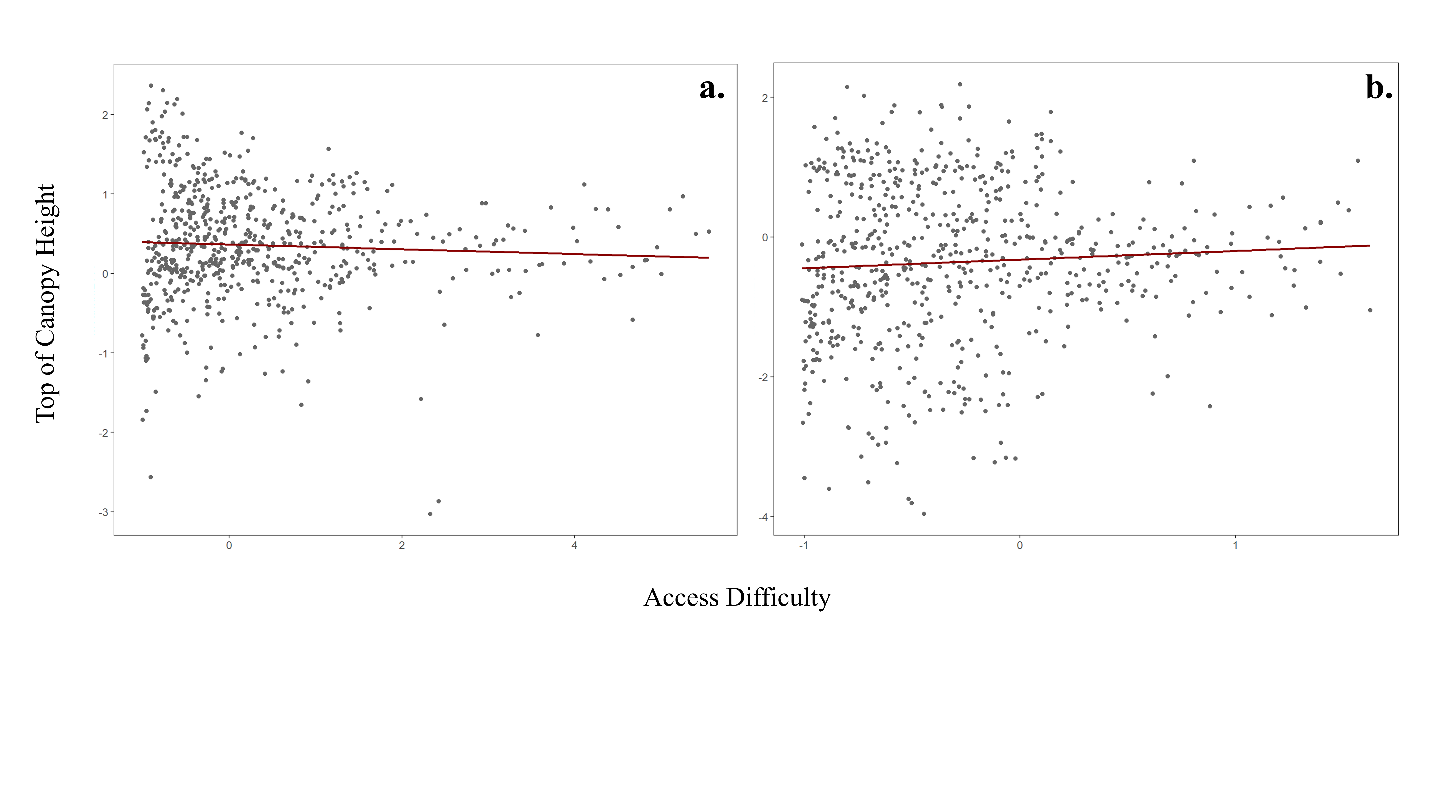

**Figure S5:** Correlation between Top of Canopy Height (TCH) and access difficulty in, (**a**) early period (r_(656)_ = -0.05, p = 0.177), and (**b**) current period (r_(656)_ = 0.06, p = 0.11). Dark red line represents regression line.
